# TERRA transcripts and promoters from telomeric and interstitial sites

**DOI:** 10.1101/2025.05.15.654290

**Authors:** Marco Santagostino, Lorenzo Sola, Eleonora Cappelletti, Francesca M. Piras, Nicolò Gennari, Marialaura Biundo, Solomon G. Nergadze, Elena Giulotto

## Abstract

The transcription of human telomeres gives rise to a family of long noncoding RNAs, named TERRA. We previously showed that TERRA transcription is driven by CpG island promoters that are composed by stretches of three types of repeats. Using the human genome assembly that was available at that time, putative promoter sequences were localized at several subtelomeres. In this work, using the T2T-CHM13v2.0 human reference genome, we found that 39 out of 46 subtelomeres contain TERRA promoters and grouped them in classes depending on their organization. We then discovered 106 intrachromosomal TERRA-like promoters, adjacent to interstitial telomeric sequences (ITSs) or far away from them. Fortyseven of these promoters are flanked and may regulate the transcription of coding genes, ncRNAs or pseudogenes. Comparative sequence analysis showed that interstitial and subtelomeric promoters belong to a previously undescribed family of segmental duplications deriving from common ancestral sequences. RT-PCR experiments in seven cell lines demonstrated that TERRA transcripts can be synthesized from ITSs. TERRA expression was always low in primary fibroblasts and HeLa cells while highly variable in the other two telomerase positive (HT1080 and HEK293) and in the three telomerase negative ALT cell lines (GM847, U2OS and VA13). The analysis of RNA-seq data from U2OS, HeLa and HEK293 cells showed that 205 ITSs were transcribed in at least one cell lines. The fraction of transcribed ITSs and the level of their transcription increased with the length of the telomeric repeat stretch. Given the large number of transcribed ITSs, we propose that these loci contribute significantly to the production of the TERRA pool.

## INTRODUCTION

Telomeres, the terminal regions of eukaryotic linear chromosomes, are chromatin structures essential for chromosome integrity. In vertebrates, the telomeric DNA is composed of extended tandem arrays of the hexanucleotide TTAGGG and is associated with the protein complex called Shelterin and with other accessory proteins (de Lange 2018). Maintenance of telomere length and protection of chromosome ends from damage is ensured by the specialized enzyme telomerase (Shay and Wright 2019), a nucleoprotein composed by a reverse transcriptase (TERT) and an RNA moiety (TERC) containing the template for telomere synthesis. The telomerase enzyme is active in the germline, in stem cells and in most tumors while, in a fraction of human cancers, a telomerase independent mechanism of telomere maintenance (Alternative Lengthening of Telomeres or ALT) is present (Azzalin 2025).

Telomeres are transcribed by RNA polymerase II into a family of long non-coding RNAs called Telomeric Repeat-containing RNA (TERRA) that, in human cells, are highly heterogeneous in length, ranging from 100 to 9000 nucleotides and are mainly composed of UUAGGG repeats (Azzalin et al. 2007). Telomeric transcripts have been identified also in mouse (Schoeftner and Blasco 2008), fish (Park et al. 2019), plants (Vrbsky et al. 2010), nematodes (Manzato et al. 2023), protozoa (Rudenko and Van der Ploeg 1989) and yeast (Luke et al. 2008). Thus, the transcription of telomeric repeats can be considered a conserved feature in eukaryotes although, in the mouse, the majority of TERRA transcripts derives from the pseudoautosomal PAR locus (Chu et al. 2017b; Viceconte et al. 2021).

Two methods were initially used to quantify TERRA molecules: northern blotting with oligonucleotide probes complementary to the telomeric repeats and reverse transcription-quantitative PCR with primers constructed on subtelomeric sequences. While the first method allowed to analyze simultaneously the molecules transcribed from all chromosome ends, with the second method TERRA molecules transcribed from a subset of specific chromosome ends could be quantified (Azzalin et al. 2007). RNA-seq experiments on purified TERRA molecules were then carried out using short Illumina reads (Porro et al. 2014a; Montero et al. 2016) or Oxford Nanopore Technology (ONT) long reads (Rodrigues et al. 2024). Recently, direct nanopore sequencing of TERRA enriched transcripts was set up (Hsieh et al. 2025). The sequencing approach, particularly with long reads, has allowed to obtain a more comprehensive picture of TERRA expression.

In humans, TERRA is transcribed from CpG island promoters that are embedded within the subtelomeric sequence immediately upstream the telomeric repeats (Nergadze et al. 2009). The pro-terminal subtelomeric regions from the long-arm of chromosomes X/Y and 10 were isolated and named TelBam3.4 and TelSau2.0, respectively (Brown et al. 1990). In that early study, it was shown that these regions comprise three minisatellites composed of stretches of 61, 29 and 37 bp repeats in centromere to telomere orientation. The 61 bp and the 29 bp repeats are separated by a sequence related to the 29 bp repeat that lies outside the array and was defined as 29like. Using a gene reporter assay, we then demonstrated that these sequences function as TERRA promoters and that the 29 bp repeat is sufficient to drive transcription (Nergadze et al. 2009). Through sequence analysis, using the GRCh37/hg19 reference genome, we identified TERRA promoters at 14 subtelomeres (1p, 2p, 3q, 4p, 5p, 8p, 9p, 11p, 15q, 16p, 17p, 19p, 21q, Xq) and on both sides of the telomere fusion at 2q14 (Nergadze et al. 2009). In the same work, we showed that the telomeric tract and the last repeat of the 37 bp unit are separated by a highly conserved sequence named pre-telomeric region or pre-tel. The regulation of TERRA transcription involves methylation of the CpG island (Nergadze et al. 2009; Le Berre et al. 2019), interaction with DNA binding proteins (Deng et al. 2012; Feretzaki et al. 2019; Vohhodina et al. 2021) and histone 3/lysine 4 methylation (Caslini et al. 2009).

Telomere transcription can increase in response to alterations of the metabolism (Diman et al. 2016) and to radiation-induced DNA damage (Al-Turki et al. 2024).

TERRA is transcribed from several chromosome ends and its expression varies between different cell lines and between different telomeres in the same cell line (Nergadze et al. 2009; Vitelli et al. 2013; Smirnova et al. 2013; Episkopou et al. 2014; Feretzaki et al. 2019; Silva et al. 2021; Savoca et al. 2023; Rodrigues et al. 2024).

TERRA is involved in the maintenance of telomere integrity contributing to the protection of chromosome ends from the activation of the DNA damage response (Porro et al. 2014b; Tutton et al. 2016; Graf et al. 2017). TERRA regulates several aspects of telomere biology, including chromatin structure (Deng et al. 2009; Arnoult et al. 2012), telomerase activity (Redon et al. 2010; Cusanelli et al. 2013) recruitment of telomeric binding proteins (Deng et al. 2009; López de Silanes et al. 2010; Scheibe et al. 2013; Abreu et al. 2022) and telomere replication (Maicher et al. 2012; Beishline et al. 2017). TERRA levels increase following induced telomeric DNA damage and, at shortened telomeres, regulating telomere length (Bettin et al. 2024; Abreu et al. 2025). The observation that deregulation of TERRA synthesis causes genomic instability and telomere loss suggested that inappropriate telomere transcription may be involved in tumorigenesis (Azzalin et al. 2007). Indeed, low levels of TERRA expression in tumors has been associated to tumor aggressiveness (Schoeftner and Blasco 2008; Deng et al. 2012; Sampl et al. 2012; Vitelli et al. 2018; Kienzl et al. 2024). Interestingly, TERRA binds not only chromosome ends but also extratelomeric genomic locations (Chu et al. 2017a; Avogaro et al. 2018). As summarized by Abreu et al (Abreu et al. 2025), the interaction of TERRA with numerous proteins involved in diverse cellular pathways suggests that it might also carry out functions unrelated to telomere maintenance (Reig-Viader et al. 2014; Wang et al. 2015; Wang and Lieberman 2016; Marión et al. 2019; Nassour et al. 2023).

Besides chromosome ends, stretches of TTAGGG repeats are present also at internal genomic sites, where they constitute the so called interstitial telomeric sequences or ITSs. ITSs have been described in many species (Meyne et al. 1990), including humans (Azzalin et al. 1997, 2001) and other primates (Ruiz-Herrera et al. 2002; Nergadze et al. 2004; Mazzoleni et al. 2017), rodents (Faravelli et al. 2002; Nergadze et al. 2007), equids (Santagostino et al. 2020) and birds (Nanda et al. 2002).

According to chromosomal localization, sequence organization and mechanism of origin, ITSs can be classified in heterochromatic, fusion and short ITSs (Ruiz-Herrera et al. 2008). Heterochromatic ITSs, extended blocks of centromeric and pericentromeric repeats, have been described in several species (Faravelli et al. 2002; Colomina et al. 2017; Clemente et al. 2020) but not in the human genome where one fusion (IJdo et al. 1991) and numerous short ITSs are present (Nergadze et al. 2007). Although short ITSs can be considered a particular type of microsatellite, they were not generated by the expansion of pre-existing units through DNA polymerase slippage (Messier et al. 1996) but were inserted in one step in the course of evolution (Nergadze et al. 2004, 2007) during the repair of DNA double strand breaks through peculiar mechanisms possibly involving the telomerase enzyme (Nergadze et al. 2004, 2007; Santagostino et al. 2020; Sola et al. 2021). Given their mechanism of origin, we may expect that insertion polymorphism of ITSs is an important source of genetic variation, as largely shown for transposable elements (Carroll et al. 2001; Witherspoon et al. 2013). However, although observed in some species, to our knowledge, it was never detected in the human population (Santagostino et al. 2020).

In the present work we characterized the organization of 39 putative subtelomeric TERRA promoters and of a family of TERRA promoter-like sequences that are positioned at interstitial sites. We then demonstrated that TERRA molecules synthesized from ITSs constitute an important component of the TERRA pool.

## RESULTS AND DISCUSSION

### Organization of subtelomeric TERRA promoters in the T2T**-**CHM13v2.0 reference genome

Taking advantage of the T2T-CHM13v2.0 genome (https://www.ncbi.nlm.nih.gov/datasets/genome/GCF_009914755.1/) (Nurk et al. 2022) we updated the analysis of human subtelomeres, completed the list of TERRA promoters and characterized their sequence organization. To test the presence of putative promoters at each subtelomere, we aligned the 61bp, 29like, 29bp, 37bp units and the pre-tel sequences from the previously described Xq TERRA promoter (Nergadze et al. 2009) (T2T-CHM13v2.0, chrX:154255311-154256622) against each 10 kb sequence immediately adjacent to the telomeric repeats of the human genome reference sequence T2T-CHM13v2.0. Sequences were then manually analyzed to verify the alignment to each repeat and to find additional sequences. With this procedure we identified, at 4 subtelomeres, a single copy sequence that we called spacer. The subtelomeres of the Y chromosome were not included in the analysis because the cell line used to generate the original T2T-CHM13v2.0 genome derived from a complete hydatidiform mole with a 46XX karyotype (Nurk et al. 2022). Supplemental Table S1 reports the genomic coordinates and sequence organization of all subtelomeres, including the ones previously described.

With this analysis we confirmed the presence of TERRA promoters at the 14 previously described subtelomeres (Nergadze et al. 2009) and identified 25 new ones. Altogether, these results suggest that, at 39 subtelomeres, TERRA promoters may drive telomere transcription. Recently, Rodrigues et al. (Rodrigues et al. 2024) analyzed the distribution of the 29 bp repeats in a T2T-derived subtelomeric reference, revealing 36 out of the 39 promoters identified here without analyzing the organization of the other repeats but positioning them within CpG rich islands.

We observed seven different sequence organizations of subtelomeric TERRA promoters (Fig. 1). Figure 1A shows a sketch of the 25 TERRA promoters characterized by the previously described sequence organization (Brown et al. 1990; Nergadze et al. 2009) with the 61-29like-29-37 repeat tracts and the pre-tel region immediately adjacent to the telomere. Fifteen of these sequences were identified in the present work (1q, 4q, 5q, 6p, 7p, 9q, 11q, 12p, 13q, 15p, 16q, 17q, 20p, 20q, 22q). At two loci (5p, 6q) we detected an additional element between the pre-tel sequence and the telomeric repeats, which we named spacer sequence (Fig. 1B). At six loci (2q, 3q, 11p, 14q, 18q and 19q) the 61-29like-29-37 repeat tracts were present while the pre-tel region was absent (Fig. 1C). At two subtelomeres (7q, 12q) the 61-29like tracts were absent and the promoter sequence comprised only the 29 and 37 repeats, the pre-tel region and the spacer sequence (Fig. 1D). In addition to these categories, three other classes were identified (Fig. 1E-G). The 29 bp repeat, which was previously shown by us to possess promoter activity in a reporter gene assay in HeLa cells (Nergadze et al. 2009), is conserved in all TERRA promoters. In a similar *in vitro* assay it was shown that the 61 bp element, which contains CTCF binding sites, favors the recruitment of RNAPII at telomeres (Deng et al. 2012). Although no data were obtained so far on the other elements found at TERRA promoters (29like, 37, pre-tel and spacer), they may function as regulators of TERRA transcription *in vivo* possibly contributing to differential expression of subtelomeres.

**Figure 1.**
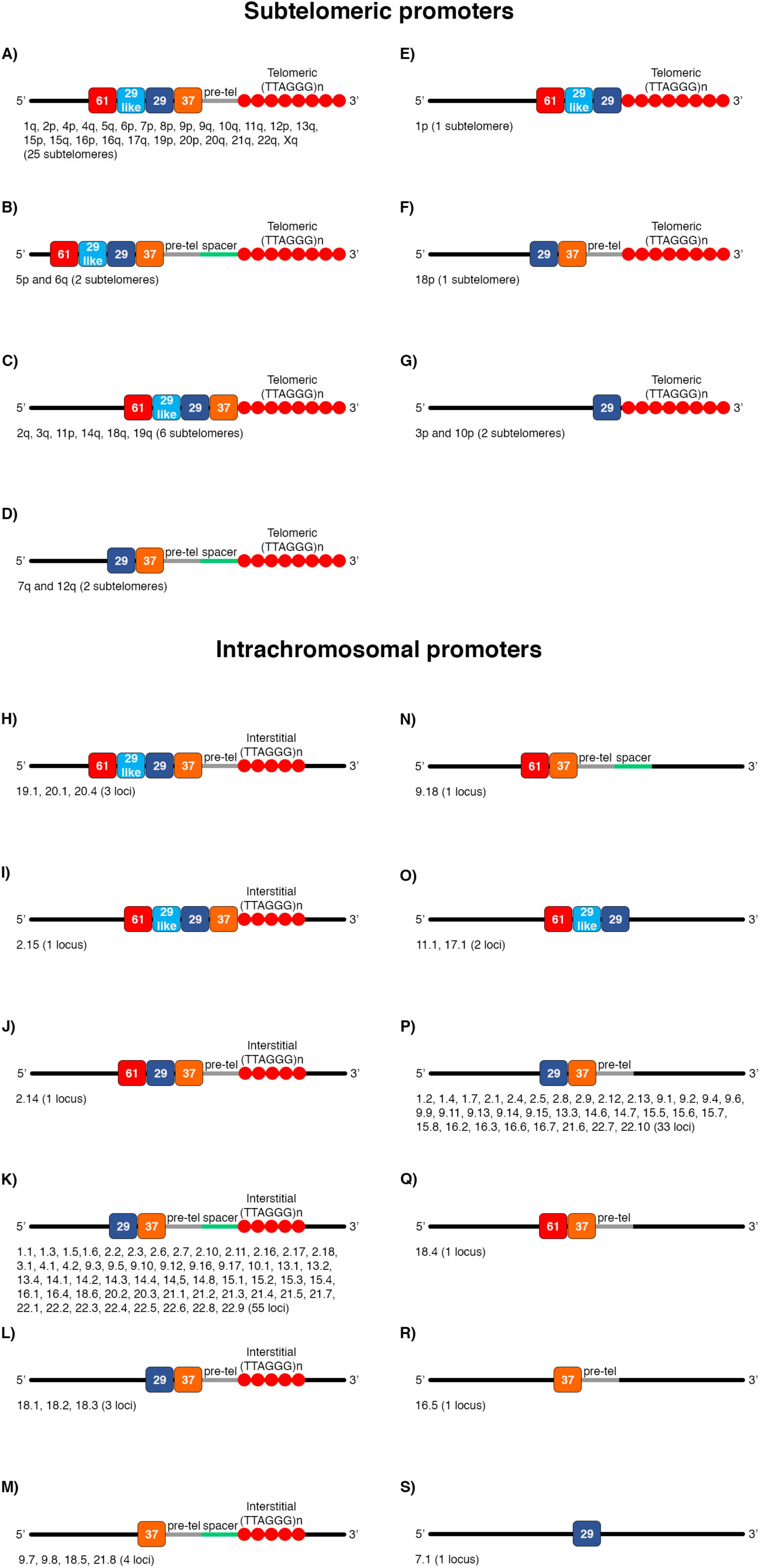
Organization of subtelomeric and interstitial TERRA promoters. Sketches of the organization of the 39 TERRA promoters found at subtelomeres (A-G), of the 67 interstitial loci flanked (H-M) and of the 39 loci not flanked by ITSs (N-S). Under each sketch, the number of loci is indicated in parenthesis. Telomeric repeats are indicated by red circles. The 61, 29like, 29 and 37 bp repeats are indicated with red, light blue, dark blue and orange rectangles, respectively. The pre-tel and the spacer sequences are indicated with gray and green lines, respectively. Flanking sequences unrelated to promoters are indicated with black lines. The subtelomeric or interstitial loci containing each type of promoter organization are listed below its sketch. Subtelomeres are indicated using the chromosome arm, interstitial loci are indicated using the names from Supplemental Table S3.

At seven subtelomeres (8q, 13p, 14p, 17p, 21p, 22p, Xp) the sequence adjacent to the telomere did not show homology to TERRA promoters (Supplemental Table S1). At these loci, the analysis of the 10 kb preceding the telomere, with a combination of different promoter predicting software (EMBOSS Cpgplot, YAPP and Promoter 2.0), did not reveal any putative promoter element with the exception of CpG islands located 4136 bp and 64 bp away from the telomeric repeats in the Xp and 14p, respectively. Sequence comparison of these subtelomeric regions extending for 5 kb from the telomere revealed that the 21p and 22p share 99% identity suggesting a common origin. The remaining five promoter-less subtelomeres did not seem to derive from a common ancestral sequence (Supplemental Figure S1). In conclusion, in the T2T-CHM13v2.0 assembly, a TERRA promoter was identified at 39 subtelomeres. At seven subtelomeres, no TERRA promoters were found suggesting that they may not be the only sequences driving telomere transcription since, as described in a following paragraph, the Xp telomere, which is not adjacent to a TERRA promoter, is transcribed in the seven cell lines tested (Figure 3A). The analysis of RNA-seq data on a subtelomeric reference sequence carried out by Rodrigues et al (Rodrigues et al. 2024) showed that these 7 subtelomeres are transcribed at different levels in the three cell lines that they tested. It would be interesting in the future to dissect these seven regions in plasmid based *in vitro* assays to characterize the putative promoters at these subtelomeres.

To test the variability of TERRA promoters in different cell lines, we compared the sequences of the subtelomeres in the T2T-CHM13v2.0 and in the GRCh38 reference genomes. In Supplemental Table S1, the results of the analysis of two genomes are reported. Twentyfive out of the 39 TERRA promoters were identified in both genome assemblies. At 5 of these informative promoters (1p, 2q, 7p, 10p and 12p) the sequence organization varied in the two assemblies. A peculiar situation was observed at the 7p subtelomere where the promoter identified in the T2T-CHM13v2.0 genome corresponded, in the GRCh38 reference, to a sequence unrelated to TERRA promoters. Interestingly, the length of the 29 bp repeat stretch, that is the minimum promoter (Nergadze et al. 2009), varied at different subtelomeres being comprised, in the T2T-CHM13v2.0 genome, between 25 and 1302 bp and corresponding to a number of repeat units comprised between about 1 and 45. The length of the other repeats varied at different loci but, at each locus, tended to be conserved in the two genomes. We cannot exclude that some of the differences between the two genomes are due to mis-assembly of the GRCh38 genome. Although reference genomes are not representative of the human population, overall, our results suggest that the sequence of some TERRA promoters may be polymorphic. The great variability of subtelomeric repeated regions in the human species supports this hypothesis (Riethman et al. 2005; Kim et al. 2025). We do not know whether sequence variability of the promoter causes TERRA expression variation among individuals, however, it may affect the efficiency of primer pairs used in RT-PCR experiments and should be taken into account when TERRA expression levels from different loci and cell lines are compared. Plasmid-based *in vitro* assays utilizing various TERRA promoter sequences were described in our original publication, in which the promoter was first identified and cloned (Nergadze et al. 2009), as well as in a subsequent study from Deng et al. (Deng et al. 2012). Although we did not detect any significant alteration of eGFP expression using plasmids with different promoter constructs, in the work of Deng colleagues (Deng et al. 2012), variation in luciferase expression was detected following deletions or mutations within the promoter sequences.

### Interstitial Telomeric Sequences (ITSs) in the T2T-CHM13v2.0 reference genome

In our previous work (Nergadze et al. 2007; Santagostino et al. 2020), we described Interstitial Telomeric Sequences in the human genome assemblies that were available at that time. Through comparative analysis with orthologous loci in non-human primates, we analyzed their origin and proposed molecular mechanisms for their insertion into genomes during DNA repair events that occurred in the course of evolution (Nergadze et al. 2004, 2007; Santagostino et al. 2020). In the present work, we updated the list of human ITSs performing a BLAST search using as query the (TTAGGG)_4_ sequence against the T2T-CHM13v2.0 reference genome. The results are reported in Supplemental Table S2 where, for each locus, the localization, length and number of mismatches per repeat is reported. While in our previous work (Santagostino et al. 2020) ITS loci were searched using as query the telomeric hexamer repeated four times with a number of mismatch/repeat ≤1, in the present work we used less stringent conditions considering as ITSs telomeric repeat loci with a minimum length of 16 bp and a number of mismatch/repeat ≤2. With this approach also degenerated loci were found. We identified 522 loci ranging in length from 16 to 1840 bp. At the majority of the ITS loci (421) the conservation of the telomeric repeats is relatively high (less than 1.4 mismatch per repeat). As previously described (IJdo et al. 1991), one of these loci, that is composed by two head-to-head stretches (HSA2q14, chr2:114027333-114028210), derives from the telomere-telomere fusion of two ancestral acrocentric chromosomes that generated human chromosome 2 (number 57 in Supplemental Table S2).

### Intrachromosomal TERRA promoters

During the search of ITS loci we discovered that 66 of them were adjacent to sequences homologous to subtelomeric TERRA promoters. At the 2q14 fusion ITS, two promoters were localized at both sites of the fusion point. Therefore, 67 TERRA-like promoters, adjacent to ITSs, were localized at intrachromosomal sites. We then detected 39 additional loci homologous to the TERRA promoter but not adjacent to telomeric repeats. Altogether, 106 interstitial TERRA promoters were found in the T2T-CHM13v2.0 genome. In Figure 1, the sequence organization (N-S) of all interstitial TERRA promoters is sketched. In Supplemental Table S3, the coordinates and the number assigned to each locus are reported.

Figure 1H shows the organization of three interstitial promoters containing the 61-29like-29-37 bp repeats and the pre-tel fragment. This is the same organization found in the majority of subtelomeres (Fig. 1A). Figures 1J and I show the organization of two promoters flanking the two ITSs, at chromosome 2q14, oriented head-to-head and deriving from the fusion of two ancestral telomeres (Ijdo et al 1991). One of these promoters is composed by the 61-29-37 bp units and the pre-tel while the other one shows the same organization of the subtelomeres shown in Figure 1C. In 55 interstitial promoters flanking an ITS (Fig. 1K) the 61 bp repeat is missing. Finally, four loci (Fig. 1M) do not contain the 29 bp core promoter.

Although at most (34 out of 39) subtelomeric promoters the 61 bp repeat is present, at most (97 out of 106) interstitial promoters this sequence is missing. It was suggested that, at telomeres, this sequence, which contains CTCF binding sites, may limit transcription proceeding towards the centromere (Deng et al. 2012). Since, as described in the next paragraph, some interstitial promoters are flanked by genes, it is tempting to speculate that the absence of the 61 bp repeat may allow transcription in both directions.

The organization of interstitial TERRA-like promoters in the T2T-CHM13v2.0 and in the GRCh38 reference genome was then compared. Only at one intrachromosomal locus (10.1 in Supplemental Table 3) the sequence organization was different in the two genomes. At the majority of loci also the length of the repeat arrays was the same in the two genomes suggesting that their sequence may be conserved in the human population.

### Comparative analysis and chromosomal localization of TERRA promoters

To test whether the sequences containing TERRA promoters are related, we downloaded the 5 kb flanking the 5’ end of each subtelomere and of each interstitial promoter. We then aligned them using the Kalign software and produced a heat map. In Supplemental Figure S2, the promoter organization, as in Figure 1, is reported. In cluster 1, 22 loci containing type K interstitial promoters flanking an ITS were included. Sixteen of these promoters were located on the acrocentric chromosomes. The majority of subtelomeric sequences were grouped in two clusters (2 and 3 in Supplemental Fig. S2) comprising 15 and 9 regions, respectively. The 5 kb regions belonging to cluster 2 were composed by a single copy sequence shared among all the 15 subtelomeres while a different single copy sequence was shared among the 9 subtelomeres in cluster 3. These single copy sequences were interrupted by a few transposons. Interestingly, four regions flanking interstitial promoters adjacent to ITSs (2.14, 2.15, 19.1 and 20.1 in Supplemental Table S3) were included in these subtelomeric clusters. The presence of subtelomeric and interstitial sites was also observed in cluster number 9. In each one of the remaining clusters (4-8 and 10) promoters sharing the same sequence organization were included.

A possible interpretation of these results is that the subtelomeric regions belonging to clusters 2 and 3, together with the interstitial regions included in the same clusters, derived from a common ancestral sequence. Successive expansions of variants that were generated through mutation may have given rise to the other clusters. In this scenario, interstitial promoters may have originated from telomeric promoters with the adjacent ITS deriving from telomeric repeats. The interstitial promoters lacking the ITS may have been generated following loss of the telomeric repeats. Alternatively, these ITS-less loci may derive from duplication events that did not involve the telomeric repeats. The second hypothesis is supported by the observation that promoters with and without ITSs are separated in different clusters (Supplemental Fig. S2). In our previous work (Nergadze et al. 2004, 2007; Santagostino et al. 2020), we showed that short ITSs were inserted in the genomes during the repair of double-strand break. Here, we identified a new class of ITSs which are adjacent to TERRA promoter-like sequences and derived from recombination events involving subtelomeric and interstitial sites.

We then localized all TERRA promoters on an ideogram of human G-banded chromosomes (Fig. 2). The intrachromosomal promoters are not randomly distributed but most of them are grouped within single chromosome bands. As shown in Supplemental Table S3, about 50% of the loci are located at less than 1 Mb from adjacent loci and, in several cases at less than 100 kb. The regions containing TERRA promoters are mainly localized near pericentromeric or subterminal regions (Fig. 2) and 29 out the 106 loci are on the short arm of the acrocentric chromosomes (13, 14, 15, 21 and 22). Some chromosomal bands on metacentric chromosomes are also particularly enriched in these loci, such as 1q21.1, 2p11.2, 9p11.2, 9q21.11, 16p11.2. In the Figure the telomeres containing TERRA promoters are also marked (red arrows). It is well known that pericentromeres, subtelomeres and acrocentric short arms are highly enriched in segmental duplications (SDs). The SD content of the human genome is about 7% and SDs make up two-thirds of acrocentric short arms (Vollger et al. 2022; Jeong et al. 2025). Given their chromosomal localization (Fig. 2) and similarity (Fig. 1 and Supplemental Fig. S2), we propose that the TERRA promoter containing regions belong to a large family of both intra- and inter-chromosomal segmental duplications that were generated during evolution via recombination events involving subtelomeric and interstitial regions. Using the SEDEF segmental duplication track from genome browser, we confirmed that 105 out of the 106 interstitial promoters are located in regions marked as duplications.

**Figure 2:**
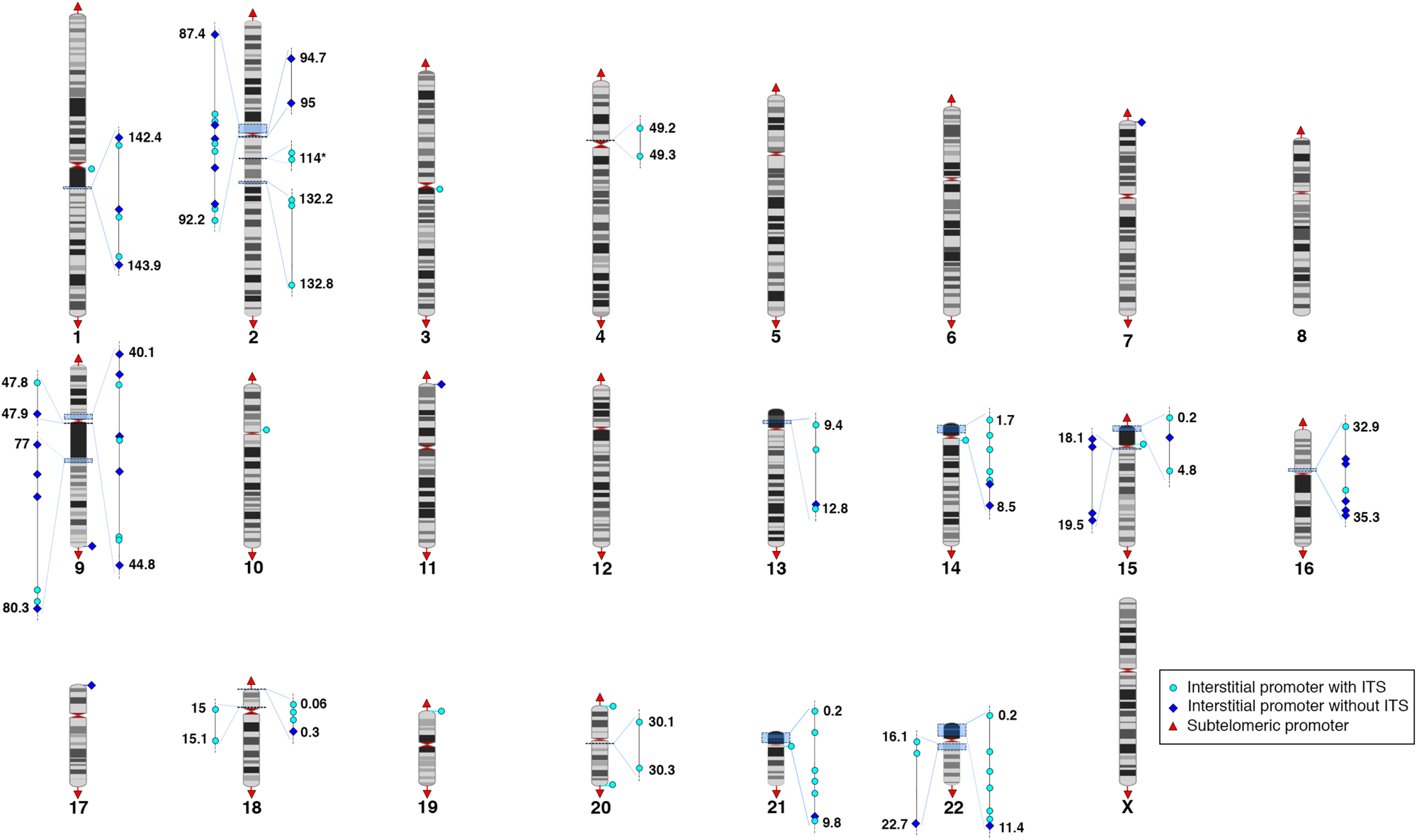
Localization of TERRA promoters in the human genome. Localization of subtelomeric (red triangles) and interstitial TERRA promoters flanked (light blue circles) or not flanked by ITSs (dark blue diamonds) on ideograms of human G-banded chromosomes. Light blue rectangles on chromosomes mark the location of groups of interstitial promoters. Magnifications on the left and right side of the chromosomes show the position of interstitial promoters in each group. Numbers at each end of the magnifications indicate the start and end coordinates of each group.

Sequence analysis of the regions containing intrachromosomal TERRA promoters was carried out using UCSC genome browser. At 47 loci, pseudogenes or genes for non-coding RNAs, whose transcription starts at less than 100 bp from a TERRA promoter, were found (Supplemental Table S4). At six loci, predicted protein coding genes were found. Genes were in the forward or in the reverse orientation relative to the promoter. At sixteen loci, genes were found on both sides of the promoter. Although localized on different chromosomes, several of these genes share high sequence identity with forward and reverse genes being included in different clusters (Supplemental Fig. S3). This is not surprising given that they are contained in duplicated regions. It is tempting to speculate that specific transcription factors may be responsible for a concerted regulation of the genes sharing common promoter organizations and that the regulation of TERRA transcription may share some of these factors.

### Transcription of TERRA molecules from interstitial telomeric sites

Recently Hsieh et al (Hsieh et al. 2025) showed that a few ITSs were transcribed in U2OS and HeLa cells. In the present work, we carried out an extensive analysis of the transcription of ITSs in different cell lines and tested the relevance of interstitial TERRA molecules, that from now on we will call iTERRA, within the total TERRA pool. First, we analyzed the transcription of some ITS loci adjacent to interstitial TERRA promoters by constructing primer pairs to amplify fragments from the pre-tel regions (Supplemental Table S5). We then tested the specificity of these primers using BLAT and *in silico* PCR in the T2T-CHM13v2.0 assembly. Five primer pairs were specific for a single locus (1.1, 2.18, 9.10, 14.3, 22.1). Given the high identity among TERRA promoter containing regions, the remaining two primer pairs amplified several loci. We also analyzed the transcription of three ITSs not flanked by TERRA promoters by constructing primer pairs on the sequence upstream the telomeric repeats, that is between 6 and 278 bp from the (TTAGGG)n array (Supplemental Table S5). These primer pairs are locus specific. As positive controls for TERRA expression, we used a primer pair specific for the 20p subtelomere, a primer pair specific for the Xp subtelomere and a primer pair that amplifies six subtelomeres (5q, 7p, 9q, 10q,13q and 16q). We measured the levels of TERRA transcription in a primary fibroblast (HPF) cell line, in three telomerase-positive (HeLa, HT1080 and HEK 293-F) and three ALT (GM847, U2OS and VA13) cancer cell lines by carrying out RT-qPCR experiments on cDNA synthesized using the ACCCTAACCCTAACC oligonucleotide. The expression levels of subtelomeres characterized by TERRA promoters was variable in the different cell lines (Fig. 3A left and middle) and were particularly high in U2OS cells but lower in the other two ALT cell lines. In primary fibroblasts, HeLa and HT1080 cells, expression levels were particularly low. These results are in agreement with previous reports where TERRA levels in primary fibroblasts, HeLa and U2OS cells were compared (Azzalin et al. 2007; Feretzaki et al. 2019; Rodrigues et al. 2024). Variable expression levels were also found at the Xp subtelomere that is characterized by the absence of a TERRA promoter. Interestingly, transcription of iTERRA molecules was detected from all the interstitial loci flanked by TERRA promoters analyzed here (Fig. 3B). As for the subtelomeric loci, great variability among cell lines was observed. The expression levels were always low in primary fibroblasts and HeLa cells while extremely variable in the other telomerase positive and in the ALT cell lines. At the ITS loci shown in the first row of Figure 3B, expression levels were higher in U2OS cells compared to HeLa and fibroblasts. However, with three primer pairs (second row in Fig. 3B), we detected higher expression levels in two telomerase positive HT1080 and HEK 293 cell lines compared to the three ALT cell lines (GM847, U2OS and VA13). The three ITS loci not flanked by a TERRA promoter were also expressed, although at relatively low levels (Fig. 3C). Since cancer cell lines are characterized by altered genomes, we tested the conservation of the ITS loci in the seven cell lines by PCR experiments on genomic DNA with the same primer pairs used for RT-qPCR. As shown in Supplemental Table S6, all genomic fragments were conserved suggesting that the variation of the detected expression levels among the cell lines is not a consequence of DNA sequence alterations.

**Figure 3:**
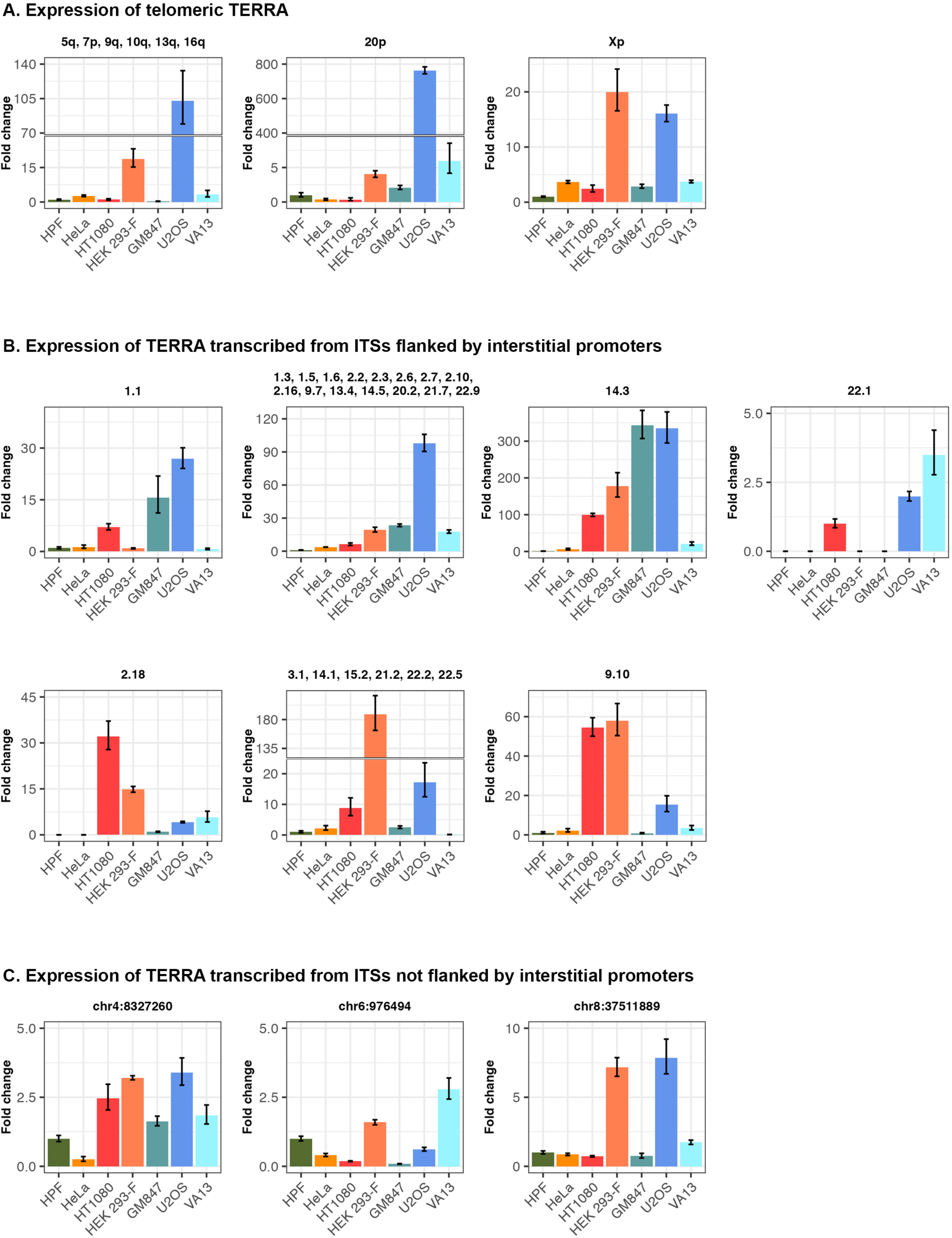
Transcription levels of iTERRA molecules measured by Real Time Quantitative PCR. Relative quantification of TERRA transcribed from telomeres (A), from ITSs flanked by TERRA promoters (B) and from ITSs not flanked by TERRA promoters (C) in a primary fibroblast (HPF) cell line, three telomerase-positive (HeLa, HT1080 and HEK 293-F) and three ALT (GM847, U2OS and VA13) cancer cell lines. The names of the loci are reported on the top of each graph. For telomeric loci the chromosome arms (A), for ITSs flanked by interstitial promoters the numbers from Supplemental Table S3 (B) and for ITSs not flanked by interstitial promoters the coordinates of the telomeric repeat starting nucleotides are reported (C). Relative quantification was calculated using the standard “2^−ΔΔCt^ method” (Livak and Schmittgen 2001). Bars indicate the minimum and maximum relative TERRA expression measured for each sample and calculated using the error propagation method. The primary fibroblasts cell line was used as reference. When no TERRA expression was detectable in fibroblasts, the cell line with the lowest expression was used as reference (HT1080 in the 22.1 locus and GM847 in the 2.18 locus).

Although only 10 random interstitial telomeric loci were tested, the results demonstrated that, in the seven cell lines, the pool of TERRA transcripts is composed by molecules synthesized from both telomeres and ITS loci.

To test whether iTERRA represents an important component of the total TERRA pool, we carried out an extensive analysis of the TERRA transcriptome. Since TERRA transcripts are poorly represented in RNA-seq datasets (Porro et al. 2014a; Rodrigues et al. 2024; Hsieh et al. 2025), we analyzed the long Nanopore sequences produced by Rodrigues and colleagues following enrichment of TERRA molecules from the human cell lines U2OS, HeLa and HEK293 (Rodrigues et al. 2024). To obtain a quantitative estimate of telomeric and interstitial TERRA molecules, we mapped the reads on the T2T-CHM13v2.0 reference genome and counted those overlapping ITSs or telomeres.

As shown in Figure 4A, U2OS cells produced the highest abundance of TERRA reads. About half of these transcripts (51%) originated from ITS containing regions. The remaining reads represented molecules transcribed from telomeres. These results suggest that, in this ALT cell line, the well described overexpression of telomere transcription may involve both types of molecules. In HeLa cells, the overall amount of TERRA transcripts is low, with the majority of them originating from ITS regions (72%). In HEK293T cells, TERRA read counts were intermediate compared to the other cell lines and, similarly to HeLa cells, most TERRA transcripts derived from interstitial loci (66%).

**Figure 4:**
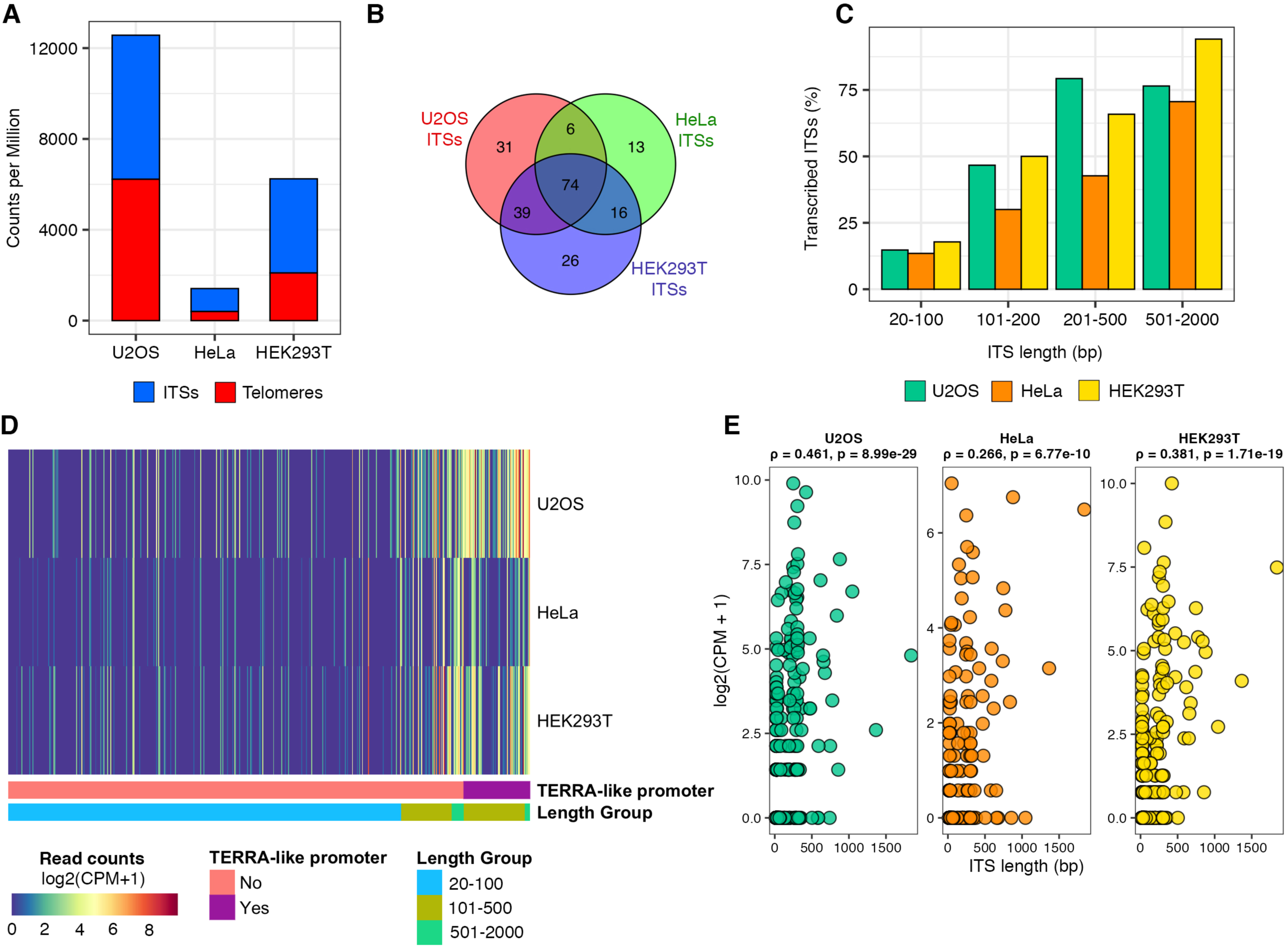
Transcription of iTERRA molecules detected in TERRA ONT-seq datasets. A) Histograms showing the abundance of TERRA reads, expressed as normalized read counts per million (CPM), in U2OS, HeLa and HEK293T cell lines. CPM for reads mapping to ITSs and extending beyond their boundaries are indicated with a blue bar, CPM for reads mapping to telomeric regions and extending into subtelomeric sequences are indicated with a red bar. B) Venn diagrams showing the number of transcribed ITS loci in HeLa (green), HEK293T (blue) and U2OS (red) cell lines. C) ITSs were grouped in four classes according to the length of the telomeric repeat stretch and the percentage of transcribed ITSs in each class was reported. The total number of ITSs in the 20-100 class was 323, in the 101-200 class was 30, in the 201-500 class was 82 and in the 501-2000 class was 17. D) Heatmap showing TERRA expression from interstitial telomeric sequences (ITSs) in U2OS, HeLa and HEK293T cell lines. Counts per million (CPM) were calculated from reads overlapping ITS genomic regions and extending beyond their boundaries. Expression values are reported as log₂(CPM + 1). ITS loci are ordered by increasing length and grouped according to ITS length and the presence or absence of TERRA-like promoters. The colored bars beneath the heatmap indicate the length group and TERRA-like promoter status, as shown in the legend. E) Scatter plots showing the correlation between ITS length (x-axis, in base pairs) and expression levels (y-axis, log₂(CPM + 1)) in the three cell lines. Spearman’s correlation coefficient and corresponding p-value are indicated.

As shown in Supplemental Table S2, the number of telomeric repeats in the genomic ITSs varies greatly. In particular, the length of the telomeric repeat tract in the 66 ITS loci flanked by a TERRA promoter is comprised between 201 and 878 bp, therefore, the length of iTERRA telomeric repeats produced by such loci is expected to be of a few hundred nucleotides while the size of promoter-less ITSs is highly variable (16-1840 bp). We then analyzed the transcripts mapping at each ITS containing region and found that 205 out of the 522 ITS loci were transcribed in at least one cell line (Figure 4B, Supplemental Table S7). The majority of these transcribed loci (66%) were shared between at least two cell lines. Interestingly, in all three cell lines, the fraction of transcribed ITSs increased with the length of the genomic telomeric repeat stretch (Figure 4C). Transcript counts varied greatly both across different ITS loci and between cell lines (Figure 4D, Supplemental Table S7). Longer ITSs, mainly associated with TERRA-like promoters, tended to show higher transcript counts compared to shorter ITSs (Figure 4D, Supplemental Table S7). The positive correlation between ITS length and read counts was statistically significant in all three cell lines (Figure 4E). This correlation may be due to expression level differences between short and long ITSs and/or to the enrichment procedure for TERRA molecules in RNA samples which may have favored the selection of longer TERRA transcripts. This bias may explain why some ITS transcripts observed by RT-qPCR were not detected in the RNA-seq datasets (Figure 3) and suggests that the fraction of transcribed ITSs may have been underestimated. Since short ITSs contain only a few telomeric repeats (Supplemental Table S2), their overall contribution to the TERRA pool remains limited not only in terms of read counts but also in the number of telomeric repeats they provide. On the other hand, transcripts originating from long ITSs or from telomeres carry greater numbers of TTAGGG repeats, and thus contribute more substantially to the total amount of telomeric sequences within TERRA containing molecules. Since TERRA-protein interactions seem to be mainly related to the formation of RNA G-quadruplex structures (Biffi et al. 2012; Liu et al. 2017; Ghosh and Singh 2020; Roach et al. 2020; Mei et al. 2021; Abreu et al. 2022), we can postulate that longer iTERRA molecules may give a greater contribution to function compared to molecules containing just a few repeats. All together, these observations suggest that iTERRA molecules constitute an important component of the total TERRA pool.

In previous work, copy number variation of TTAGGG repeats, was detected at ITS loci (Mondello et al. 2000; Aksenova and Mirkin 2019; Santagostino et al. 2020). Therefore, we expect that, in the genome of the cell lines analyzed here, the length of several ITSs might be slightly different from the T2T-CHM13v2.0 reference genome. However, according to the mechanism of microsatellite expansion and contraction, that is DNA polymerase slippage (Messier et al. 1996), ITS length variation should not affect significantly our analysis that is based on the ITS length in the T2T-CHM13v2.0 reference genome (Figure 4C-E).

## CONCLUDING REMARKS

In conclusion, in this work, we presented a detailed description and comparative sequence analysis of the 39 subtelomeric TERRA promoters and of the 7 promoter-less subtelomeres in the T2T-CHM13v2.0 reference genome. We then discovered that 106 TERRA promoters are localized at intrachromosomal sites and, together with the subtelomeric promoters, belong to a new family of segmental duplications. Therefore, the recombination mechanisms that gave rise to the duplicons were also responsible for the insertion in the genome of this new class of ITSs. After updating the list of ITS in the T2T-CHM13.v2 reference genome, we obtained the most relevant finding: in all the cell lines analyzed, an important component of TERRA is transcribed from interstitial telomeric repeats.

Important questions remain open on the function of TERRA, iTERRA and interstitial TERRA promoters. The observation of high iTERRA levels in U2OS cells where telomeric TERRA is also hyper-expressed suggests that the two types of transcripts may be coregulated possibly through binding to common factors. To this regard, we previously showed by ChIP-seq that the human telomeric repeat factors 1 and 2 (TRF1 and TRF2), which are part of the Shelterin complex, bind to a subset of ITSs in the BJ-HELTRas tumor cell line (Simonet et al. 2011). This result was confirmed by Yang et al (Yang et al. 2011) together with the observation that, in the HTC75 fibrosarcoma cell line, RAP1, another shelterin protein, binds to a few interstitial telomeric sites. Later, Mukherjee at al (Mukherjee et al. 2019) observed that, in the HT1080 fibrosarcoma cell line, thousands of extra-telomeric TRF2 binding sites were enriched in potential G-quadruplex structures. Interestingly, we found that 82 of these TRF2 peaks map on ITS loci, with 31 of them coinciding with peaks identified by Simonet et al (data not shown). In light of these observations, it is tempting to postulate that some of the putative TRF2 extra-telomeric functions may be mediated by ITS binding. In this scenario, since depletion of TRF2 is known to cause TERRA upregulation (Porro et al. 2014a), it would be interesting to test whether the expression of iTERRA is also increased.

It is known that TERRA associates with telomeres forming R-loops and that RAD51 recombinase is required for this association at chromosome ends suggesting that TERRA may contribute to homology directed recombination mechanisms such as those occurring in ALT cells (In et al. 2025). An intriguing hypothesis is that iTERRA may also play a role in these pathways through invasion of telomeric repeat tracts in *cis* or in *trans*.

Since subtelomeric sequences undergo frequent rearrangement, telomeric transcripts from different cell lines may not be correctly mapped on specific chromosome ends of the T2T-CHM13.v2 reference genome. In addition, as mentioned above, the procedure for telomeric transcripts enrichment may have led to the selection of longer repeat arrays. The availability of RNA-seq data from T2T isogenic reference genomes and of RNA sequencing procedures avoiding any type of pre-enrichment will allow a more precise analysis and quantification of TERRA and iTERRA transcripts.

## MATERIALS AND METHODS

### Identification of subtelomeric and interstitial TERRA promoters

Subtelomeric TERRA promoters were identified in the T2T-CHM13v2.0 genome assembly (https://www.ncbi.nlm.nih.gov/datasets/genome/GCF_009914755.1/) (Nurk et al. 2022). We downloaded the 10 kb sequences immediately adjacent to each telomere in the T2T-CHM13v2.0 genome assembly using UCSC Genome Browser (https://genome.ucsc.edu). Using Multalin, we searched each subtelomere for sequences related to the previously described Xq TERRA promoter (chrX:154255311-154256622, T2T-CHM13v2.0 assembly) (Nergadze et al. 2009). Sequences were manually analyzed to define the organization of the promoters. Using Clustal Omega (Sievers et al. 2011) sequence identity with the Xq reference was determined and varied between 80 and 100% for the 61 bp, between 64 and 99 % for the 29like, between 55 and 100% for the 29 bp, between 71 and 100% for the 37 bp and between 65 and 100% for the pre-tel. Subtelomeres that did not contain sequences related to the TERRA promoter were analyzed for the presence of predicted promoters using CpGplot (https://www.ebi.ac.uk/Tools/seqstats/ emboss_cpgplot/), YAPP (http://www.bioinformatics.org/yapp/cgi-bin/yapp.cgi/), and Promoter 2.0 (https://services.healthtech.dtu.dk/services/Promoter-2.0/).

Interstitial TERRA promoters were BLAST-searched (https://blast.ncbi.nlm.nih.gov/Blast.cgi) in the T2T-CHM13v2.0 genome assembly using the sequence chrX:154255311-154256622 as query. The search was performed using this setup: blastn algorithm, max target sequences = 100; automatically adjust parameters for short input sequences = disabled; expect threshold = 10; word size = 11; max matches in a query range = 0; match/mismatch scores = 2, -3; gap costs = existence: 5, extension: 2, and filters for low complexity regions and species-specific repeats disabled. The automatic adjustment of search parameters for short sequences was disabled.

Further manipulations of the hit list were carried out using tools available on the Galaxy platform (https://usegalaxy.org/) (Afgan et al. 2016). Sequences were analyzed manually and through Multalin alignment against the chrX:154255311-154256622 sequence to analyze promoter sequences.

### Comparative analysis of TERRA promoters and flanking genes

To perform sequence comparison, we downloaded the 5 kb flanking the 5’ end of each subtelomeric and of each interstitial promoter. Sequences were aligned using the Kalign software (Lassmann and Sonnhammer 2005) (https://www.ebi.ac.uk/jdispatcher/msa/kalign?stype=dna) to produce a percentage identity matrix that was converted into a heatmap using the R package "pheatmap" (Kolde 2018) (https://rdrr.io/cran/pheatmap/). Using the UCSC genome browser, we searched in the T2T-CHM13v2.0 genome for the presence of putative or annotated genes whose transcription starts within 100 bp from each interstitial promoter. The genomic sequences corresponding to these gene were downloaded and aligned using the Kalign software to produce a percentage identity matrix that was converted into a heatmap.

### Localization of TERRA promoters on the human chromosomes

The kariogram of human G-banded chromosomes showing the localization of the subtelomeric and interstitial TERRA promoters was prepared using the R package RIdeogram (Hao et al. 2020) and cytoband information from the T2T-CHM13v2.0 genome assembly available at the UCSC Genome Browser.

To test whether regions containing intrachromosomal TERRA promoters belong to a family of Segmental Duplications, we used the Table Browser tool to find the promoters that have any overlap with regions in the SEDEF Segmental Duplications (https://genome.ucsc.edu/cgi-bin/hgTrackUi?hgsid=2516619039_nMpaYeBSA3J2vYcU3JP5EXXUa27D&db=hub_3671779_hs1&c=chr9&g=hub_3671779sedefSegDups).

### Search for ITS loci in the human T2T-CHM13v2.0 genome assembly

To update the list of human ITSs, the sequence (TTAGGG)_4_ was used as query for a BLAST search (https://blast.ncbi.nlm.nih.gov/Blast.cgi/)(Altschul et al. 1990) against the human genome reference sequence *Homo sapiens* T2T-CHM13v2.0 (https://www.ncbi.nlm.nih.gov/datasets/genome/GCF_009914755.1/)(Nurk et al. 2022). The search was performed with the “blastn” algorithm and the following parameters: Max target sequences = 100, Expected threshold = 10, Word sizes =11, the automatic adjustment parameters for short sequences, low complexity regions and species-specific repeats regions were disabled. Further manipulations of the hit list were carried out using tools available on the Galaxy platform (https://usegalaxy.org/) (Afgan et al. 2016).

The exact position and organization of each ITS in the human genome was manually determined using the UCSC Genome Browser (https://genome.ucsc.edu/) (Kent 2002).

### Cell culture

Seven human cell lines were used to test TERRA transcription at interstitial loci: human normal primary fibroblasts (HPFs), three telomerase-positive (HeLa, HEK293-F adenovirus-transformed human embryonic kidney and HT1080 fibrosarcoma cells) and three ALT cancer cell lines (U2OS osteosarcoma cells, VA13 SV40-immortalized lung fibroblast and GM847 SV40-immortalized skin fibroblasts cells). All cell lines were cultured in 5% CO_2_ at 37°C. Human primary fibroblasts were propagated in Dulbecco’s Modified Eagle’s Media (DMEM, Euroclone), supplemented with 20% Fetal bovine serum (Euroclone), 1% L-glutamine (Sigma), 2% non-essential amino acids (Euroclone) and 1% Penicillin-Streptomycin (Sigma). Cancer cell lines were cultured in DMEM supplemented with 10% Fetal bovine serum (Euroclone), 1% L-glutamine (Sigma), 1% non-essential amino acids (Euroclone) and 1% Penicillin-Streptomycin (Sigma).

### RNA preparation, cDNA synthesis and RT-qPCR

Total RNA was extracted from 10^7^ cells using QIAzol Lysis Reagent (QIagen) according to the manufacturer’s protocol. To remove genomic DNA, we repeated the following procedure twice: RNase-free DNase I (Thermo Scientific) (1 U per µg of RNA) at 37°C for 30 min; "on column" DNase I digestion and purification using the Zymo Research RNA Clean & Concentrator kit (Zymo Research) according to the manufacturer’s instructions.

To synthesize the cDNA for quantitative Real Time PCR experiments, we set up reverse transcription reactions in a 20 µl volume containing 5 µg of total RNA, 10 pmoles of the ACCCTAACCCTAACC oligonucleotide to reverse transcribe TERRA, 5 pmoles of the AGTTGCCAGCCCTCCTAGA oligonucleotide to reverse transcribe the reference gene β2-microglobulin. Reverse transcriptions were carried out at 58°C using the Maxima H-Reverse Transcriptase (Thermo Scientific) according to the manufacturer’s protocol for GC-rich transcripts. Finally, to prevent the formation of RNA:DNA hybrids, the reaction was incubated with 1 U of RNase H (Promega) at 37°C for 30 min. RT-reactions were prepared using the same protocol and substituting the Maxima H-Reverse Transcriptase with an equivalent volume of water.

Forward and reverse primers for RT-qPCR reactions were designed on the DNA sequence upstream the TTAGGG array of telomeres and of ITS loci, within 400 bp from of the TTAGGG repeats (Supplemental Table S5). Quantitative Real Time PCRs were performed using the 2X GoTaq qPCR master mix (Promega) in a 96-well reaction plate (Thermo Scientific). For each primer pair, 3 cDNAs, a no-template control and two non-retrotranscribed controls were tested from each cell line. Mixes were prepared according to the manufacturer’s protocol in a total volume of 20 µl containing 2 µl of cDNA, and the forward and reverse primer (5 pmoles each) (Supplemental Table S5). Reactions were carried out using the PikoReal Real-Time PCR System (Thermo Scientific). Cycling parameters included an initial denaturation at 95°C for 7 min, followed by 45 cycles composed by the following steps: 95°C for 10 sec, annealing coupled to extension at 62°C for 30 sec with fluorescence detection. A final fluorescence acquisition for melting curve analysis (60°C to 95°C) was done.

The relative TERRA expression was calculated according to the standard “2^−ΔΔCt^ method” (Livak and Schmittgen 2001). Data were normalized against the hB2M gene using the human primary fibroblast cell line as reference. When no TERRA expression was detectable in fibroblasts, we used the cell line with the lowest expression as reference. For each cell line, the minimum and maximum relative TERRA expression detected for each primer pair in each triplet of reactions was calculated using the error propagation.

### Analysis of TERRA ONT-seq datasets

TERRA ONT-seq datasets from U2OS (SRR25394385), HeLa (SRR25394384), and HEK293T (SRR25394383) cell lines produced by Rodrigues et al. (Rodrigues et al. 2024) were retrieved from the NCBI SRA Archive. Adapters were trimmed using Porechop (Galaxy Version 0.2.4+galaxy1) (Wick et al. 2017). Reads were mapped to the human T2T-CHM13v2.0 reference genome using minimap2 (Galaxy Version 2.28+galaxy1) (Wick et al. 2017) in long-read splice mode (-x splice). Primary alignments were selected with Samtools (version 1.15.1) (Li 2018). Normalized counts (CPM) of reads overlapping ITS loci or telomeres were obtained using the pysam module (https://github.com/pysam-developers/pysam). Reads overlapping these regions by at least 12 bp and extending beyond the ITS or telomeric boundaries were included in the analysis. Reads that fully mapped telomeric regions, without aligning to the adjacent reference sequences, were counted as telomeric. Plots were generated using the pheatmap, ggplot2 and VennDiagram R packages. Statistical significance of the correlation between read counts and ITS length was assessed using Spearman’s correlation test in R.

### Genomic DNA preparation and PCR

Genomic DNA from the seven cell lines was purified using phenol/chloroform standard protocols. PCR experiments on genomic DNA were carried out with the same primer pairs used for RT-qPCR (Supplemental Table S6). PCR reactions were performed in a 25 µL final volume with 100 ng of genomic DNA, 15 pmoles of each primer, 0.2 mM dNTP, 1X Green Buffer (Promega) and 0.5 units of GoTaq DNA polymerase (Promega). An initial denaturation step was carried out at 95°C for 2 min and 30 sec, followed by three amplification cycles at 95°C for 1 min, annealing temperature (Supplemental Table S6) for 1 min and 72°C for 45 sec. The first three cycles were followed by 32 additional cycles: 95°C for 40 sec, annealing temperature for 40 sec, 72°C for 40 sec. Final extension was at 72°C for 5 min. The size of the PCR products was checked by electrophoresis in 2% agarose gel.

## ACKNOWLEDGMENTS

We would like to thank Claus Azzalin and Joana Rodrigues (Istituto de Medicina Molecular Joao Lobo Antunes, Universidade de Lisboa, Lisbon) for their valuable insights during the revision of this manuscript. We also thank Francesco Lescai (Department of Biology and Biotechnology, University of Pavia) for his helpful suggestions on the bioinformatic analyses.

